# Fear Generalization as Threat Prediction: Adaptive Changes in Facial Exploration Strategies revealed by Fixation-Pattern Similarity Analysis

**DOI:** 10.1101/125682

**Authors:** Lea Kampermann, Niklas Wilming, Arjen Alink, Christian Büchel, Selim Onat

## Abstract

Animals can effortlessly adapt their behavior by generalizing from past experiences, and avoid harm in novel aversive situations. In our current understanding, the perceptual similarity between learning and generalization samples is viewed as one major factor driving aversive generalization. Alternatively, the threat-prediction account proposes that perceptual similarity should lead to generalization to the extent it predicts harmful outcomes. We tested these views using a two-dimensional perceptual continuum of faces. During learning, one face is conditioned to predict a harmful event, whereas the most dissimilar face stays neutral; introducing an adversity gradient defined only along one dimension. Learning changed the way how humans sampled information during viewing of faces. These occurred specifically along the adversity gradient leading to an increased dissimilarity of eye-movement patterns along the threat-related dimension. This provides evidence for the threat-prediction account of generalization, which conceives perceptual factors to be relevant to the extent they predict harmful outcomes.

## Introduction

Generalization is a fundamental cognitive ability, making it possible to use previously acquired knowledge in novel situations (Pavlov, 1927; Hovland, 1937; Spence, 1937; Guttman and Kalish, 1956; Shepard, 1987; Tenenbaum and Griffiths, 2001; Struyf et al., 2015). For instance, a previously unseen food item that is potentially harmful can be avoided, if it resembles one that is known to be harmful. In potentially threatening situations, this ability is called fear (or equivalently aversive) generalization and provides an important benefit to the organism by shaping behavior adaptively (Dunsmoor et al., 2011; Greenberg et al., 2013; Onat and Büchel, 2015; Resnik and Paz, 2015; Kahnt and Tobler, 2016).

According to one prominent view, the perceptual model, it is the perceptual similarity between previously learned and novel generalization samples that determines the degree of generalization. In this view, the degree to which a novel stimulus is considered as harmful is directly related to the degree of overlap between shared sensory features with a previously learnt harmful stimulus. An alternative view proposes that fear generalization is an active cognitive act (Shepard, 1987), related specifically to the prediction of potential threat in uncertain situations (Onat and Büchel, 2015). According to the threat-prediction model, perceptual factors can contribute to fear generalization, but only to the extent they are predictive of harmful events (Lashley and Wade, 1946; Shepard, 1987; Tenenbaum and Griffiths, 2001; Baddeley et al., 2007; Soto et al., 2014). In majority of previous studies, perceptual similarity between the generalization and learning samples has been explicitly used as a cue for signaling the threat. As a result, threat-prediction has commonly been confounded by perceptual similarity, making it impossible to dissociate their independent contributions. Therefore, it remains to be shown whether threat-prediction or perceptual similarity explains fear generalization best in a paradigm that dissociates perceptual similarity and threat prediction.

In order to dissociate independent contributions of perceptual similarity and threat-prediction, we investigated fear generalization using a two-dimensional perceptual space (Fig. 1A) with faces arranged along a circle within this space (Butter, 1963; Onat and Büchel, 2015). By pairing one item with an aversive outcome and keeping the most dissimilar one neutral (opposite face separated by 180 degrees), we introduced an adversity gradient defined exclusively along one perceptual dimension. The perceptual space can therefore be decomposed into threat-*specific* and *unspecific* components (Fig 1B, middle panel), where the latter models perceptual similarity independent of adversity. Hence, using this set of stimuli made it possible to dissociate independent contributions of perceptual factors related to similarity as such, from those relevant for the prediction of threat.

**Fig 1.**
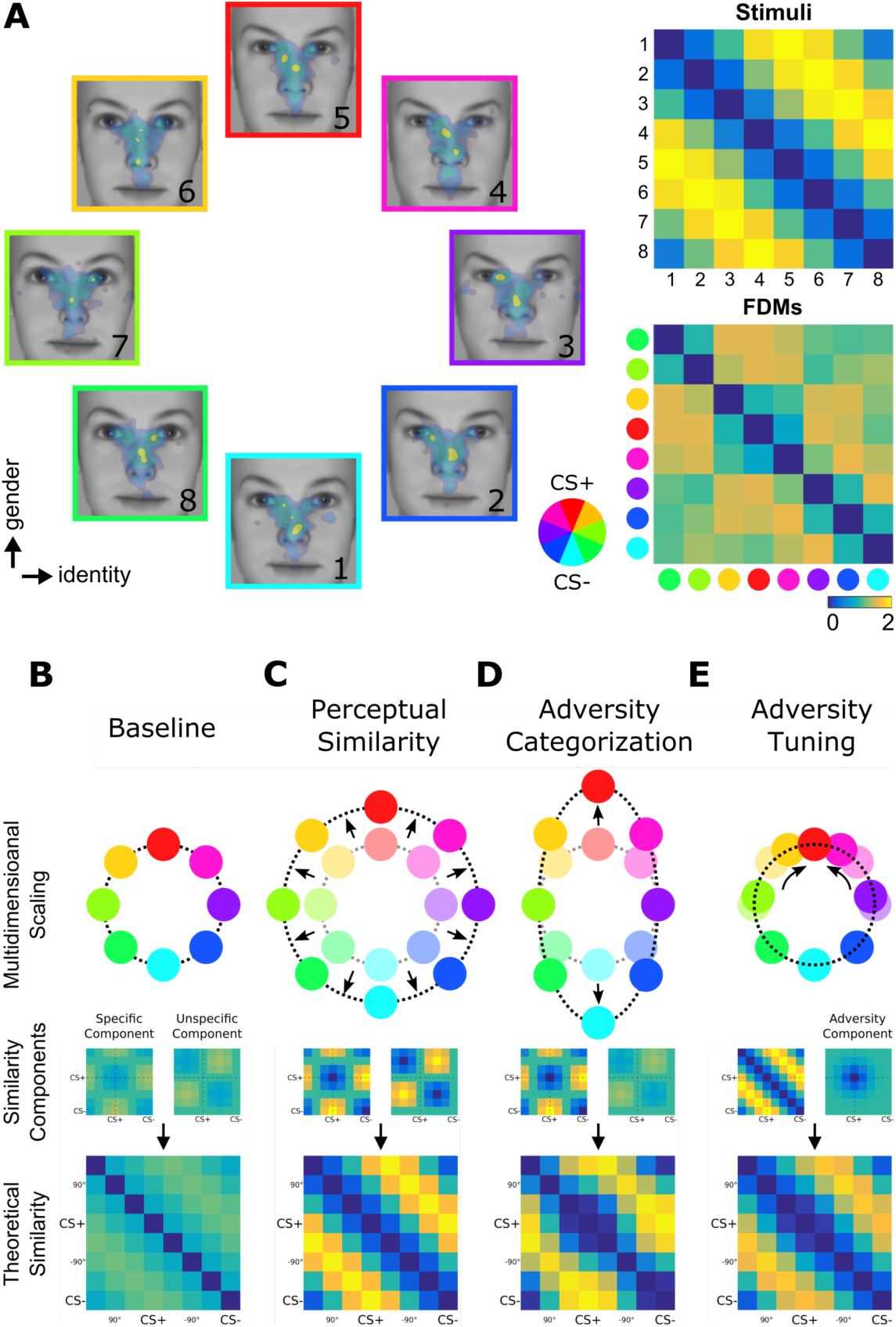
Fixation-Pattern Similarity Analysis. **(A)** 8 exploration patterns (colored frames) from a representative individual overlaid on 8 face stimuli (1 to 8) calibrated to span a circular similarity continuum across gender and identity dimensions. A pair of maximally dissimilar faces was randomly selected as CS+ (red) and CS– (cyan; see color wheel). Similarity between the 8 faces was calibrated to have a perfect circular organization with lowest dissimilarity (blue) between neighbors, and highest dissimilarity (yellow) for opposing pairs. FPSA summarizes the similarity relationship between the 8 exploration patterns as a symmetric 8×8 matrix (bottom right panel). 4th and 8th columns (and rows) are aligned with the CS+ and CS–, respectively. **(B-E)** Multidimensional scaling representation of four theoretical similarity relationships investigated with FPSA (top row). Each colored node represents one exploration pattern (red: CS+; cyan: CS–), where internode distances are proportional their dissimilarity (bottom row). Shaded nodes in **(C-E)** depict the pre-learning state in **(B)**. Dissimilarity matrices are further decomposed onto basic similarity components (middle row) centered either on the CS+/CS– (specific component) or +90°/–90° faces (unspecific component). A third component shown in **(E)** is uniquely centered on the CS+ face (adversity component). In **(B)**, equal contribution of basic components results in circularly similar exploration patterns. In **(C)**, a stronger equal contribution results in a better global separation of all exploration patterns (denoted by radial arrows). In **(D)**, a stronger contribution of the specific component results in a biased separation of exploration patterns specifically along the adversity gradient defined between the CS+ and CS– nodes. In **(E)**, the adversity component centered on the CS+ face can specifically decrease the dissimilarity of exploration patterns for faces similar to the CS+, resulting in circularly shifted nodes (circular arrows) while preserving the global circularity of the similarity relationships.

We based our analysis on multivariate eye-movement patterns. This is motivated by the fact that faces on the stimulus continuum are characterized by subtle differences. Humans explore such complex stimuli serially (Itti and Koch, 2001; Parkhurst et al., 2002; Tatler et al., 2005; Onat et al., 2014) and rely heavily on their overt attentional resources due to well documented perceptual bottleneck for forming detailed representations of complex scenes (Treisman and Gelade, 1980; Hochstein and Ahissar, 2002; Ahissar and Hochstein, 2004). Therefore, eye-movement recordings can faithfully mirror the strategies used for foraging information from different faces, which in turn inform us on the way how aversive learning affects cognitive strategies along the specific and unspecific directions (Dowd et al., 2016; König et al., 2016; Henderson and Hayes, 2018).

For the analysis of eye-movement patterns, we introduced a similarity-based multivariate method that we termed fixation-pattern similarity analysis (FPSA). Instead of focusing on how often arbitrary parts of a face are fixated (Malcolm et al., 2008; Schurgin et al., 2014; Hessels et al., 2016), FPSA uses simultaneously all available fixations to derive a similarity metric. One great benefit of this approach was that the circular organization of our stimuli allowed us to formulate hypotheses on how the similarity relationships between exploration patterns could change along the threat-specific and -unspecific directions of the perceptual continuum. This way, perceptual similarity and threat-prediction account translate to different weighting of specific and unspecific components (Fig 1B-E, middle panels).

First, the perceptual similarity account predicts that exploration patterns should enable evaluation of perceptual similarity with the CS+ face. This would result in exploration strategies that strongly mirror the physical similarity relationships between faces (Fig 1C) and lead potentially to globally increased dissimilarity following learning. As similarity information varies both on the specific as well as the unspecific dimensions, the *perceptual similarity* hypothesis predicts a stronger, but more importantly equal contribution of the underlying specific and unspecific components (Fig 1C, middle panel). In contrast, the threat-prediction account requires that eye-movements support a categorization process for faces based on their outcome as harmful vs. safe (Shepard, 1987; Ohl et al., 2001; Vervoort et al., 2014; Dunsmoor and Murphy, 2015; Qu et al., 2016). To achieve this, exploration strategies would be tailored to target locations that are maximally discriminative of the CS+ and CS– faces. This would lead to exploration patterns becoming more similar for faces sharing similar features with the CS+ and CS– faces, while simultaneously predicting an increased dissimilarity between these two sets of exploration patterns (Fig 1D). Increased similarity only along the threat-relevant dimension would then result in an ellipsoid representation of similarity relationships. Therefore, the *adversity categorization* hypothesis would lead to an increase of the adversity specific component without influencing the unspecific component (Fig 1D, middle panel). Alternatively, but still in line with the threat-prediction framework, exploration strategies could support threat-prediction by tailoring viewing patterns to quickly identify the CS+. A new sensorimotor strategy exclusively for the adversity predicting face would lead to a localized change in the similarity relationships around the adversity-predicting CS+ face (Fig 1E). Thus, the *adversity tuning* hypothesis predicts an increased similarity of exploration patterns for faces closely neighboring the CS+ face, decaying proportionally with increasing dissimilarity to the CS+ face. This strategy is not exclusive and does not directly map to the specific or unspecific components.

## Results

We created 8 face stimuli that were organized along a circular similarity continuum characterized by subtle physical differences in facial elements across two dimensions (gender and identity; see SFig 1 for stimuli). We calibrated the degree of similarity between faces using a simple model of the primary visual cortex known to mirror human similarity judgments (Yue et al., 2012) (see SFig 2 for calibration). The physical similarity relationship between all pair-wise faces conformed with a circular organization (Fig 1A, top right panel), such that dissimilarity varied with angular difference between faces (lowest for left and right neighbors and highest for opposing faces) with equidistant angular steps. Participants (n = 74) freely viewed these faces before and after an aversive learning procedure (Fig 2A) while we measured their eye-movements. During the conditioning phase, one of the eight faces was introduced as the CS+, being partially reinforced with an aversive outcome (UCS, mild electric shock in ∼30% trials). The CS– was the face most dissimilar to the CS+ (separated by 180°) and was not reinforced. During the subsequent generalization phase, all faces were presented again and the CS+ continued to be partially reinforced to prevent extinction of the previously learnt association. These reinforced trials were excluded from the analysis. To ensure comparable arousal states between the baseline and generalization phases, we administered UCSs also during the baseline period, however they were fully predictable as their occurrence was indicated by a shock symbol (Fig 2A). Furthermore, we inserted null trials during all phases (i.e. trials without face presentation but otherwise exactly the same) in order to obtain reliable baseline levels for skin-conductance responses.

**Fig 2.**
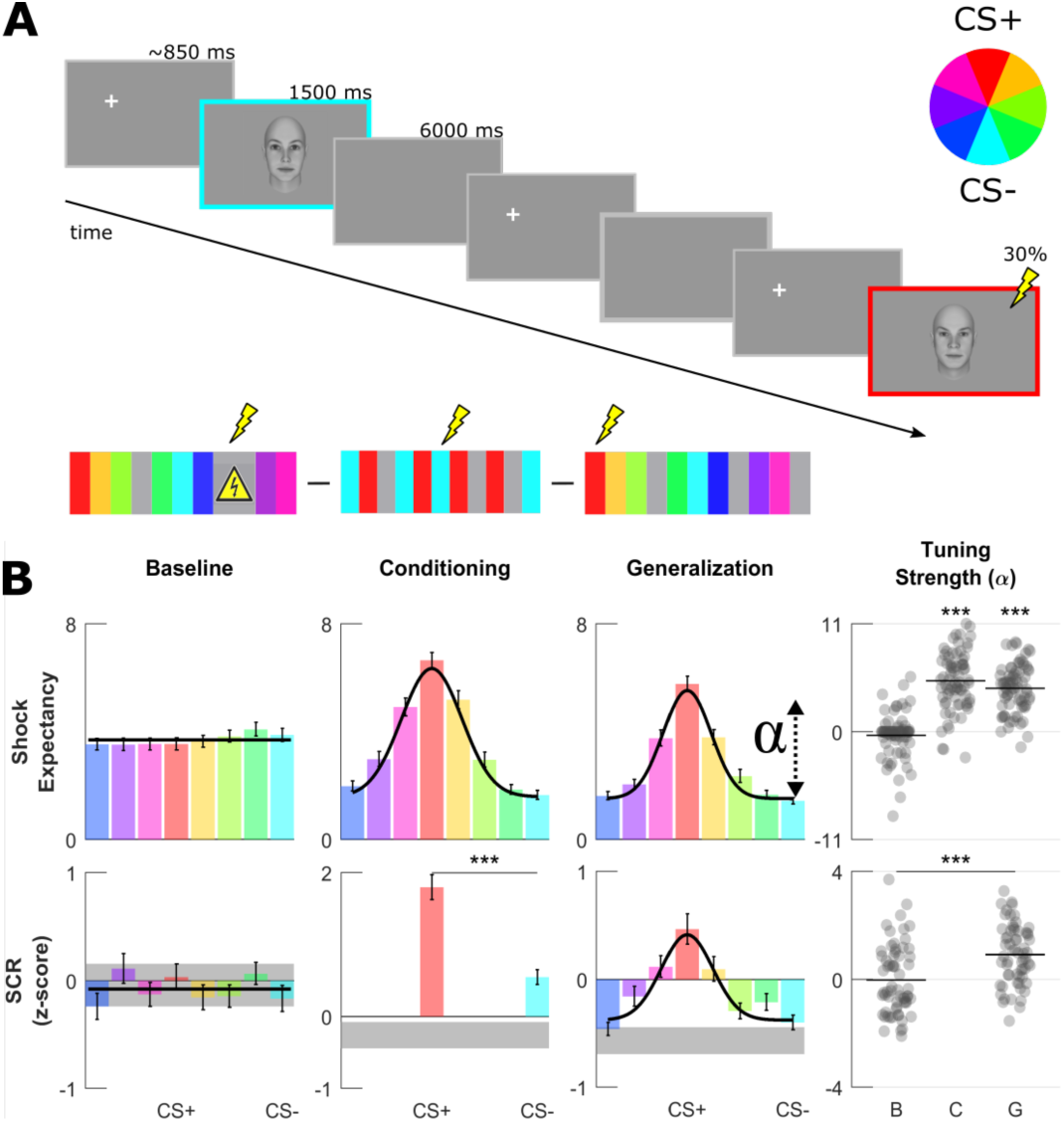
Aversive learning procedure and univariate generalization profiles. **(A)** On every trial, one randomly selected face was presented for 1.5 seconds preceded by a fixation cross placed outside of the face on either the left or right side. In null trials, no face was shown (*gray frame*) resulting in a SOA of ∼6 or ∼12s. For each volunteer, a pair of most dissimilar faces was randomly selected as the CS+ (*red*) and CS– (*cyan*, see color wheel). During baseline, UCSs (shock sign) were completely predictable by the presentation of a triangular signboard. During conditioning and generalization, the CS+ face was paired with an aversive outcome in ∼30% of trials allowing recording of responses from non-reinforced CS+ trials. **(B)** Group-level fear-tuning based on subjective ratings of UCS expectancy (n = 74) and SCR (n = 63) for different phases. Responses are aligned to the CS+ of each volunteer separately (errorbars: SEM across subjects). Black horizontal lines or curves indicate the winning model (p < .001, log-likelihood ratio test), i.e. horizontal null model or the Gaussian model. Gray shaded areas in SCR depict response amplitudes evoked by null trials (mean and 95% CI). Scatter plots show amplitude parameter of Gaussian fits (denoted by alpha symbol) for each volunteer. Horizontal lines within the scatterplots depict group-level means, asterisks indicate significant differences in α (compared to baseline phase, paired t-test, ***: p < .001).

### Fear tuning profiles in subjective ratings and autonomic activity

As expected, the effect of learning was mirrored both in autonomous nervous system activity as well as subjective ratings of UCS expectancy (Fig. 2B first and second rows). During the conditioning phase, skin-conductance responses (SCR) were on average 3.8 times higher for CS+ than CS– (Fig 2B, middle panel bottom row, paired t-test, p < .001) indicating that the CS+ face gained a stronger aversive quality already during the conditioning phase. In line with this view, expectancy ratings gathered right at the end of the conditioning phase were also highest for the CS+ face (Fig 2B, middle panel top row).

As in previous studies, we characterized fear generalization by computing fear tuning profiles based on subjective ratings and SCR. In both recording modalities, responses decayed with increasing dissimilarity to the CS+ face and reached minimal values for CS–. We modeled these with a Gaussian function centered on the CS+ face. At the group-level, model comparison favored the flat null model over the Gaussian in both recording modalities before learning (p = .44 for SCR, p = .17 for ratings, log-likelihood ratio test; black horizontal lines in Fig 2B). However, following the conditioning phase the Gaussian model fitted the data significantly better (comparison to flat null model, p < .001 for SCR and subjective ratings, log-likelihood ratio test). Fear-tuning profiles at the single-subject level were in agreement with the overall group-level picture. We summarized fear-tuning profiles of individual participants with the amplitude parameter of the fitted Gaussian function, which characterizes the modulation depth of fear tuning (i.e. the strength of fear tuning) after accounting for baseline shifts. The average amplitude parameter following the conditioning phase was significantly bigger than the baseline phase in both recording modalities (paired t-test, p < .001, see Fig 2B). In summary, univariate fear-tuning in SCR and subjective ratings confirmed that aversive learning was successfully established and transferred towards other perceptually similar stimuli.

### Multivariate fear tuning profiles in eye movements

We analyzed exploration behavior using fixation density maps (FDMs). These are two-dimensional histograms of fixation counts across space, smoothed and normalized to have unit sum (see methods section for details). When computed separately for different faces, FDMs indicate how much different parts of a given face receive attentional resources. For each volunteer, we computed a dissimilarity matrix that summarized all pairwise comparisons of FDMs (using 1 - Pearson correlation as a pattern distance measure) and averaged these after separately aligning them to each volunteer’s custom CS+ face (shown always at the 4th column and row in Fig 3A). Furthermore, in order to gather an intuitive understanding of learning-induced changes in the similarity geometry, we used multidimensional scaling (jointly computed on the 16×16 matrices, i.e. all pairwise combinations of 8 conditions from baseline and generalization phase). Multidimensional scaling (MDS) summarizes similarity matrices by transforming observed dissimilarities as closely as possible onto distances between different nodes (Fig 3B) representing different viewing patterns in 2D, therefore making it easily understandable at a descriptive level.

**Fig 3.**
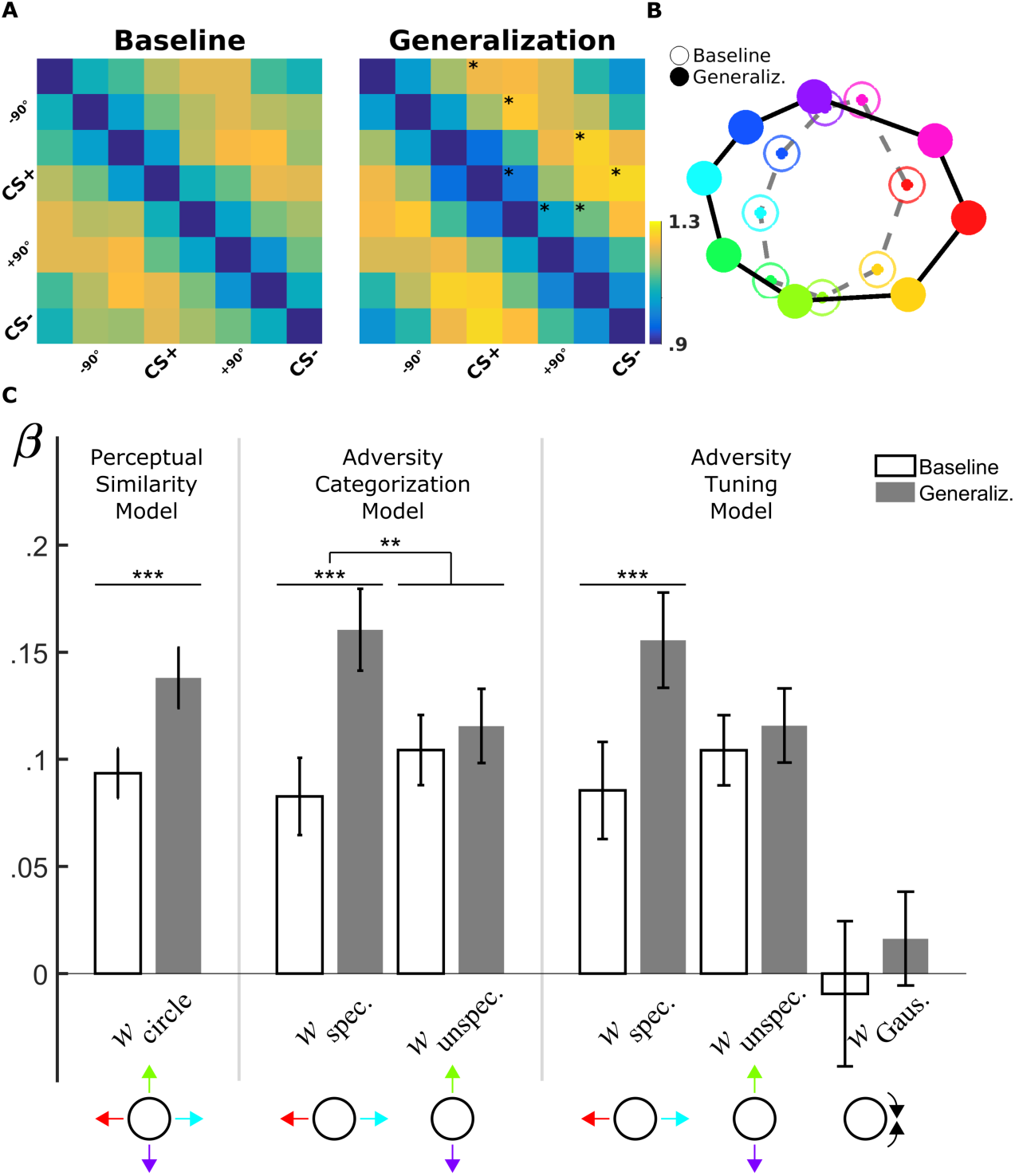
Fixation-Pattern Similarity Analysis. **(A)** Dissimilarity matrices of exploration patterns for baseline (*left panel*) and generalization phases (*right panel*). Fourth and eight columns (and rows) are aligned with each volunteer’s CS+ and CS– faces, respectively. Asterisks on the upper diagonal denote significant differences from the corresponding element in baseline. **(B)** Two-dimensional multidimensional scaling method used for visualization of the 16×16 dissimilarity matrix (not shown) comprising baseline and generalization phases. Distances between nodes are proportional to the dissimilarity between corresponding FDMs (*open circles*: baseline; *filled circles*: generalization phase; same color scheme as in Fig 1). **(C)** Bar plots (M ± SEM) depict predictor weights estimated for single-participants before (white bars) and after (gray bars) aversive learning (*Left:* Bottom-up Saliency; *Middle:* Adversity Categorization; *Right:* Adversity Tuning model). *w*_circle_: weight for the circular component, which is the sum of equally weighted specific and unspecific components; *w*_specific_/*w*_unspecific_: weights for specific and unspecific components; *w*_Gauss_: weight for adversity component centered uniquely on the CS+. (**: p < .01; ***: p < .001, paired *t-test*).

Already during the baseline period the dissimilarity matrix was highly structured (Fig 3A). In agreement with a circular similarity geometry and the MDS depiction, lowest dissimilarity values (1.05 ± .01; M ± SEM) were found between FDMs of neighboring faces (i.e. first off-diagonal), whereas FDMs for opposing faces separated by 180° exhibited significantly higher dissimilarity values (1.21 ± .01; paired t-test, *t*(73) = 7.41, p < .001). Using the Perceptual Model, we investigated the contribution of physical characteristics of the stimulus set to the observed pre-learning dissimilarity structure (Fig 1B). This model uses a theoretically circular similarity matrix (consisting of equally weighted sums of specific and unspecific components) as a linear predictor. This model performed significantly better compared to a null model consisting of a constant similarity for all pairwise FDMs comparisons (for Perceptual Model adjusted *r*^2^ = .09; log-likelihood-ratio test for the alternative null model: p < 10-^5^; BIC_NullModel_ = −96.1, BIC_Perceptual_ = 244.7; see S1 Table for the results of model fitting). We additionally fitted the Perceptual Model for every volunteer separately (Fig 3C). Model parameters at the aggregate level were significantly different from zero (Fig 3C; *w* _Circle_ = .09 ± 0.01, M ± SEM; *t*(73) = 7.99, p < .001) indicating that exploration strategies prior to learning mirrored the physical similarity structure of the stimulus set. This provides evidence that fixation selection strategies are, at least to some extent, guided by physical stimulus properties during viewing of neutral faces.

We observed significant changes between baseline and generalization dissimilarity values in an element-wise comparison (Fig 3A, indicated by asterisks). This provides evidence for learning-induced changes in the similarity relationships. Following learning, the same Perceptual Model was again significant (adjusted *r*^2^ = .35; *p* < .001, log-likelihood ratio test), but now performed better compared to the baseline phase (BIC_Perceptual_ = 244.7 for the baseline vs. BIC_Perceptual_ = 1697.4 for the generalization phase; see S2 Table for model fitting results). Critically, we found a significant increase in the model parameter from baseline to generalization phase (*w* Circle = 0.13 ± 0.01; paired *t*-test, *t*(73) = 4.03, *p* < .001; Fig 3C compare two leftmost bars) suggesting a global increase in dissimilarity between FDMs. Overall, these results are compatible with the view that aversive learning led to a better separation of exploration patterns globally, in agreement with the Perceptual Model (Fig 1C), which predicted a major contribution of perceptual similarity on exploration patterns following learning.

However, according to the multi-dimensional scaling method the separation between exploration patterns occurred mainly along the adversity gradient defined by the CS+ and CS– faces, whereas the separation along the orthogonal unspecific direction did not exhibit any noticeable changes (Fig 3B). We thus extended the circular Perceptual Model to capture independent variance along the two orthogonal directions using the Adversity Categorization model (Fig 1D). Model comparison indicated that the more flexible model performed better than the simpler Perceptual Model (BIC_Perceptual_ = 1697.4 vs. BICAdversityCateg. = 1957.5; adjusted *r*^2^ = .48 vs. 0.35; see STable 3-4 for fitting results with the Adversity Categorization model on baseline and generalization phases, respectively). A two-factor (experimental phase × predictor type) repeated measures ANOVA showed a significant main effect of experimental phase (i.e. before vs. after conditioning, F(1, 73) = 6.29, p < .001), as well as a significant interaction of predictor (specific vs. unspecific) with experimental phase F(1,73) = 8.53, p < .005). Accordingly, post-hoc analyses showed that while the unspecific component did not change from baseline to generalization phase (t(73) = .75, p = .45), the specific component gained stronger contribution (t(73) = 4.64, p < .001), and these learning-induced changes were significantly larger in the specific as compared to unspecific component (*t*(73) = 2.92, p < .005, paired t-test). This observation provides evidence that increased overall dissimilarity was driven by changes in the scanning behavior specifically along the task-relevant adversity direction.

We next evaluated whether the observed anisotropy in the similarity geometry was relevant for the aversive quality associated with the CS+ face. We tested the hypothesis that stronger anisotropy (wspecific – wunspecific) could predict stronger aversive learning, as measured with the modulation depth of fear-tuning profiles coming from subjective ratings and skin-conductance responses. We found weak, but significant evidence for an association with an increased tuning strength in the ratings (r = .25, p = .03), which was only marginally present for SCR (r = .23, p = .07). This suggests that separation of exploration patterns along the adversity gradient is related to aversive learning, and rules out the possibility that it could simply result from increased exposure to the CS+ and CS– faces throughout the conditioning period.

The remodeling of the similarity geometry along the adversity gradient can also be accompanied by exploration strategies that are specifically tailored for the adversity predicting face but not for CS– resulting in localized changes only around the CS+ face. We subjected this view to model comparison by augmenting the previous model with a similarity component that consisted of a two-dimensional Gaussian centered on the CS+ face. Positive contribution of this predictor would lead to more similar exploration patterns specifically around the CS+ (Fig 1E). It can thus capture changes in similarity relationships that are specific to the CS+ face. The model comparison procedure favored the simpler Adversity Categorization model over the augmented Adversity Tuning model (BIC_AdversityCateg_. = 1957.5 vs. BIC_AdversityTuning_.= 1923.6 during the generalization phase; adjusted *r*^2^ = .48; see S5-6 Table for fitting results with Adversity Tuning model in baseline and generalization phases, respectively). Hence the increase in the number of predictors did not result in a significant reduction in explained variance. In line with this result, the parameter estimates for the CS+ centered adversity component were not significantly different from zero neither in baseline or generalization phases (w_Gaussian_ = .009 ± .034 in baseline, *t*(73) = -0.27, p = 0.78; w_Gaussian_ = 0.02 ± 0.02 in generalization, *t*(73) = .74, p = 0.46; Fig 3C). Also, pair-wise differences between parameter estimates did not reach significance (t(73) = 0.72, p = 0.47). We therefore conclude that further extensions of the Adversity Categorization model to include components for adversity-specific changes around the CS+, did not result in a better understanding of the adversity-induced changes in the similarity geometry of exploration strategies.

### Temporal and spatial unfolding of adversity specific exploration

While SCRs and subjective ratings provide insights about the aggregate cognitive evaluations of a given stimulus, eye-movements have the potential to provide information on how these cognitive evaluations unfold over both spatial and temporal domains. We therefore repeated the FPSA using eye-movement data that originated from different temporal or spatial windows. For this analysis, we focused on participants that were able to correctly identify the CS+ based on their shock expectancy ratings (CS+ > CS–, n = 61). First, to gain insights on the temporal dynamics of specific exploration, we used a moving window of 500 ms with steps of 50 ms and repeated FPSA using the Adversity Categorization Model (Fig. 4A). Before learning, the time-course of adversity-specific and unspecific components was not distinguishable. However, during the subsequent generalization phase they diverged rather early. We tested the anisotropy difference in similarity geometry (w_spefic_ – w_unspecific_) before and after learning. The difference reached significance first at the time window corresponding to the interval 400-900 ms after stimulus onset (Fig. 4A *top row*, paired t-test, p = 0.03). As humans explore visual scenes serially with fixational eye-movements, the order of fixations (1st fixation, 2nd fixation, and so on) is another natural metric to evaluate the temporal progress (Tatler et al., 2005). The same analysis indicated that adversity-specific exploration started following the first fixation (note that the first fixation is the landing fixation on the face following stimulus onset; Fig. 4A) and stayed constant during the stimulus presentation. Overall, the temporal FPSA indicated that humans started to forage for adversity-specific information early, as soon as after the first landing fixation.

**Fig 4.**
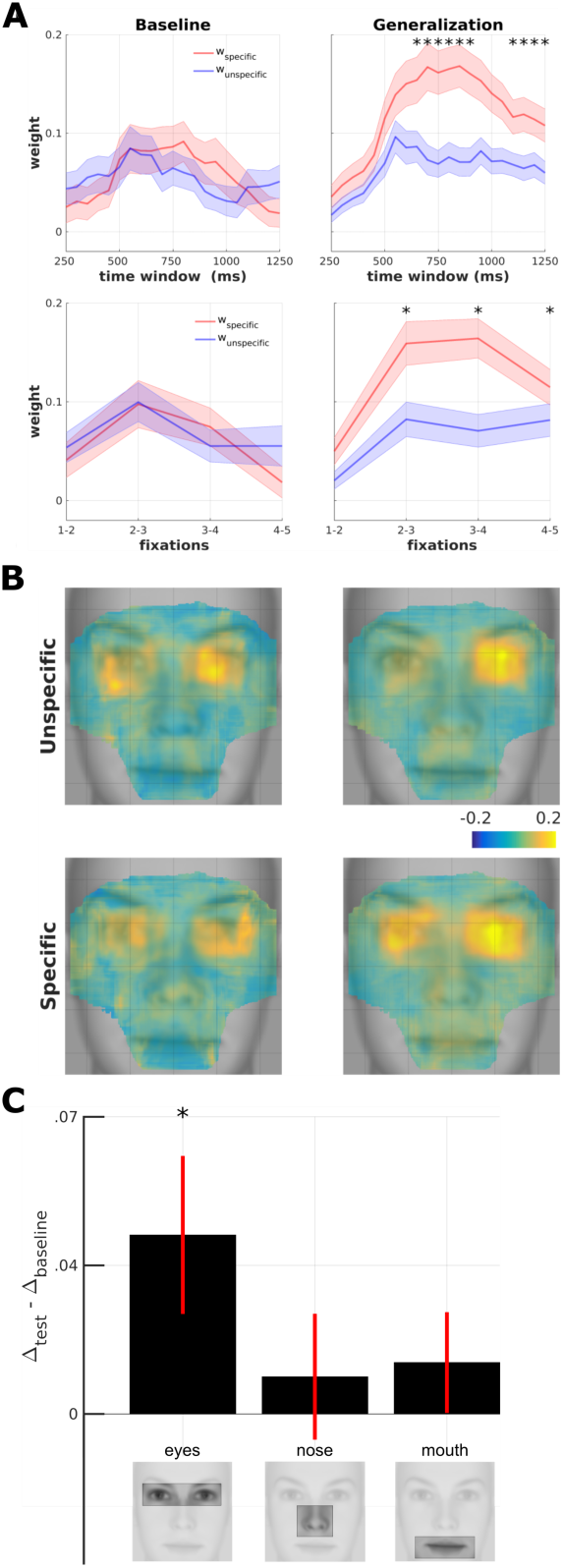
Spatio-temporal Fixation-Pattern Similarity Analysis. **(A)** Temporal development of adversity-specific and unspecific exploration strategies (n = 61). Parameters of the adversity categorization model (*red*: specific; *blue*: unspecific) are computed using a moving time window of 500 ms at intervals of 50ms for baseline (*left panels*) and test phase (*right panel*). Numbers on the x-axis denote the center of the moving window. Second row depicts the same analysis with fixation points sorted by rank. Asterisks indicate statistically significant interaction testing for difference in anisotropy between test vs. baseline (*: p < .05; Shaded area: SEM) **(B)** Four maps resulting from the searchlight-FPSA on FDMs from before and after conditioning (*left vs. right columns*) and for unspecific and specific (*top vs. bottom rows*) model parameters overlaid on an average face. The map is masked to contain 90% of all fixation density. **(C)** Difference of anisoptropy (w_specific_ – w_unspecific_) between before and after aversive learning (*test – baseline*) for three different ROIs (*: p < 0.01, paired t-test).

One limitation of FPSA is that the information about the spatial origin of specific or unspecific exploration strategies is lost. To circumvent this limitation, we ran FPSA at localized portions of the FDMs in a similar manner to a searchlight analysis in brain imaging (Kriegeskorte et al., 2007). For a given spatial portion (defined by a square window of 30 pixels, ∼1 visual degree), we fitted the Adversity Categorization Model, and assigned specific and unspecific weights to the center position of the searchlight (Fig. 4B). The map that is obtained by repeating this analysis at all spatial locations provides an indication of the facial locations that are explored either with a specific or unspecific exploration strategy. We found that both before and after learning, specific and unspecific components were strongly localized around the eye region. We tested the difference in anisotropy between the baseline and test phases within the three commonly used regions of interest at different facial elements (eyes, nose and mouth; ROIs shown in Fig 4C) (Walker-Smith et al., 1977; Malcolm et al., 2008; Schurgin et al., 2014; Hessels et al., 2016). We found this interaction to be significant only in the eye region (Fig. 4C; paired t-test uncorrected, p < .024). This result could be explained by an increased number of fixations around the eye region following learning. This could potentially result in a stronger signal-to-noise ratio for the estimation of different similarity components (Diedrichsen et al., 2011). Indeed, the eye region in this study accounted of ∼80% of all fixations (SFig. 3, *top row*). However, during the generalization phase, we observed an increase in the weight of the specific components around the eye region, despite a decrease in the overall fixation density (∼5%, *top row* in SFig. 3). Overall, searchlight FPSA indicated that adversity-specific exploration strategies were specifically tailored to find differences around the eye region.

### Comparison of FPSA to ROI-based analysis of eye-movements

In order to control whether the multivariate approach of FPSA on eye movement patterns exceeded the sensitivity of common ROI-based analyses on fixation counts, we computed changes in fixation counts in the three common regions of face stimuli, i.e. eyes, nose and mouth (same as depicted Fig 4C). If conditioning before the generalization phase lead to an increased saliency of facial features that are diagnostic of the CS+ face, one would expect a non-flat fear tuning in the number of fixations towards these facial features, which would receive more fixations with increasing similarity to the CS+ face. In line with previous reports (Walker-Smith et al., 1977), eyes together with the nose region were the most salient locations across the baseline and generalization phases, and attracted ∼84% of all fixation density, whereas the mouth region had only a marginal contribution with ∼3.5%. Investigating changes from baseline to generalization phase, we found that aversive learning increased the number of fixations directed at the nose (+4.4%) and mouth (+0.8%) regions at the expense of the eye region (=.3%). Most importantly, model comparison on fixation density favored the flat null model for all regions even at low statistical thresholds in all facial elements across both baseline and generalization phases (p > 0.05, log-likelihood test; SFig. 3). While a weak tuning was apparent in the mouth region during generalization, this did not reach significance (p > 0.1). Therefore, our observations at the group-level were limited to unspecific changes between phases that were independent of the adversity gradient introduced through conditioning.

## Discussion

The present work tested the validity of competing hypotheses on aversive generalization. We used a stimulus continuum that was defined by perceptual similarity, as well as threat-related information as two independent perceptual factors. As an extension of similarity-based multivariate pattern analyses used to investigate representational content of neuronal activity in fMRI (Kriegeskorte et al., 2008) and MEG/EEG (Cichy et al., 2014; Kietzmann et al., 2017), we introduced FPSA to characterize learning-induced changes in eye-movement behavior during free viewing of faces. As expected, before aversive learning exploration patterns mirrored the inherent circularity of stimulus similarity. This is compatible with the view that the exploration of neutral faces in our experimental settings is, at least to some extent, guided by physical characteristics, in contrast to a completely holistic viewing strategy (Peterson and Eckstein, 2012). Aversive learning led to a specific increase in dissimilarity along the direction of the adversity gradient. This adversity-specific exploration strategy appeared early, as soon as following the landing fixation, and lasted continuously for the duration of stimulus presentation, mainly to forage perceptual evidence around the eye region. This shows that behavior during fear generalization was specifically tailored to detect differences along the threat-relevant stimulus dimension. Overall, these changes in exploration patterns indicate that fear generalization can be understood as an active process related to the prediction of threat, and not simply a response to perceptual similarity between learned and novel samples. Perceptual similarity plays a decisive role to the extent it predicts the occurrence of relevant events.

Our results showed that the separation between the CS+ and CS– exploration patterns were increased the most. There are at least two different scenarios, which could lead to this observation (Fig. 5A). This regards the way how learning potentiates sensory features as being diagnostic for the prediction of harmful outcomes, and making them target locations for attentional allocation during active viewing. In the first scenario, aversive learning potentiates unique combination of visual features that are specific to the harm-predicting item, namely the CS+ face identity (Fig. 5A, *top panel*). Alternatively, aversive learning can lead to an increased saliency for discriminative features that separate best the harm- and safety-predicting prototypes. In this view, aversive learning consists of recovering the vector of features that defines the adversity gradient (Fig. 5A, bottom panel). This feature vector could overlap with a categorical information that is either naturally present (such as gender, ethnicity or emotional expression), or learned *de novo* with experience (Kietzmann and König, 2010; Qu et al., 2016). Both scenarios lead to an increased separation between the CS+ and CS– poles as observed in this study. However, they have divergent predictions when tested with stimuli organized in three concentric circles (Fig. 5B). If learning modifies uniquely identity-specific representations, faces from outer and inner circles that are close to the CS+ would be explored similarly, hence resulting in a shrinkage of the similarity geometry around the CS+ face. On the other hand, if learning is based on a vector representation, the pattern separation would add to differences that are already present, resulting in three concentric ellipses centered on the same point. Future investigations can distinguish these two hypotheses by testing whether exploration patterns during generalization results in a shift in the center of similarity geometry towards the CS+ face or not.

**Figure 5.**
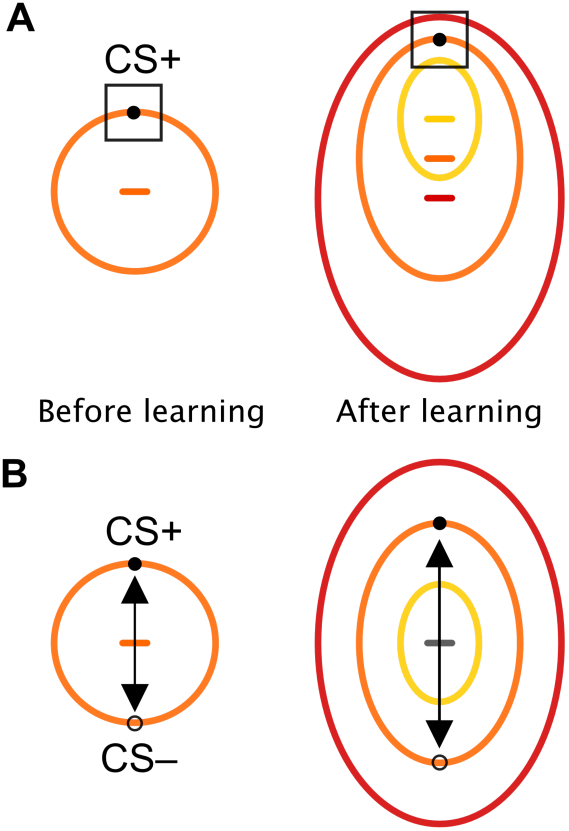
Predictions on the influence of aversive learning on sensory-motor foraging strategies. Before learning, exploration strategies follow the circularity of the stimulus space (*orange curve*) in line with their physical characteristics. **(A)** During conditioning, humans learn the feature vector (*black arrow*) that separates best harmful and safety predicting stimuli. When tested subsequently with stimuli organized as three concentric stimulus gradients, this scenario predicts three concentric ellipses sharing the same center of gravity (*gray horizontal line*) **(B)** In another scenario, during conditioning humans learn the specific feature values that predict an harmful outcome (*black square*). When tested with the concentric stimulus set, this scenario predicts faces that are similar to the CS+ face to be explored similarly. This would result in a global shift in the center of gravity towards the CS+ face (*yellow, orange and red horizontal lines*).

Our results contrast with previous conceptualizations of fear generalization as driven simply by perceptual similarity. According to the perceptual model, fear generalization is viewed as resulting from perceptual similarity between current observations and an event that is known to truly predict a behaviorally relevant outcome. This model has received substantial support from the associative learning theory, which provided a mechanistic framework for the perceptual model (Lissek et al., 2014). In this view, the extent to which a novel stimulus evokes fear-related responses is directly proportional to the overlap in the associative connections formed previously during learning. Different theories within this associative framework have emphasized either elemental (Rescorla and Wagner, 1972) or configural (Pearce, 1987, 2002) factors in the computation of a perceptual similarity metric for the guidance of generalization. However, these invariably reduced the cognitive ability for generalization to a byproduct of perceptual similarity between learning and generalization samples, as has been indicated in recent reviews (Soto et al., 2014; Dunsmoor and Murphy, 2015). Our observation of a specific separation of exploration patterns is not reconcilable with the simple perceptual model, as this model predicts an equal contribution of all perceptual factors for the prediction of behaviorally relevant events. Instead, a specific separation of exploration patterns indicates that additional mechanisms are required for a selective weighting of perceptual factors related to the prediction of behaviorally relevant outcomes.

Selective changes in the similarity of exploration patterns is compatible with the view that following learning, the nervous system achieves a categorization process along the smooth perceptual continuum (Ohl et al., 2001; Visser et al., 2013; Dunsmoor et al., 2014; Qu et al., 2016). Such a categorization could lead onto selective changes in exploration patterns, which would in turn lead to an increased efficiency of information transmission downstream. Notably, theoretical work has already showed that generalization can be viewed as the formation of an internal model of the environment in order to predict the occurrence of behaviorally relevant events. Shepard’s generalization theory conceives generalization as the formation of a binary categorical zone along a smooth perceptual continuum for the prediction of harmful or safe events (Shepard, 1987). In this view, empirical generalization profiles can be understood as resulting from different stimuli to be registered into either the one or the other category in a probabilistic manner. In the same line, Bayesian frameworks refer explicitly to probability distributions to represent internal beliefs for the formation of generalization strategies (Tenenbaum and Griffiths, 2001; Tenenbaum et al., 2006). These theoretical works have already pointed to the possibility that humans can adaptively tailor generalization strategies by appropriately structuring their internal beliefs even in situations which are not defined by perceptual factors. Our results provide supporting evidence that these mechanisms have their validity even in settings that are defined solely by perceptual factors.

Introduction of the FPSA methodology was not a choice, but rather a necessity to test the outlined competing hypotheses. First, eye-movement patterns during free-viewing of faces are characterized by strong inter-individual variability (Walker-Smith et al., 1977; Mehoudar et al., 2014; Kanan et al., 2015; Coutrot et al., 2016). This was also true in the present study, where we could train linear classifiers to predict a volunteer’s identity with an average accuracy of ∼80% based on fixation maps (results not shown). This is possibly one major reason why we did not observe adversity-tuned generalization profiles at different facial ROIs (SFig. 3). While this inter-individual variability in exploration patterns can dilute the sensitivity of ROI-based approaches, FPSA requires instead a consistent change in the similarity relationships of exploration patterns. Therefore, one major benefit of FPSA was making it possible to test the validity of hypotheses in the presence of highly variable inter-subject eye-movement patterns. Second, and equally importantly, it is not clear how one could test different hypotheses outlined here using one-dimensional generalization profiles — collected from subjective ratings, autonomous recordings or fixation counts from individual ROIs. Therefore, another benefit of FPSA was making it possible to subject a rich set of hypotheses to statistical testing.

Could our results be explained by an unbalanced exposure to the CS+ and CS– faces that were presented during the conditioning phase? As participants have seen these faces more often, one can argue that this could potentially bias eye movement patterns. While we cannot completely exclude a contribution of exposure, we have shown that anisotropy correlates with indicators of aversive learning as measured by subjective ratings and, to lesser extent with autonomous activity. Furthermore, unpublished observations on neuronal recordings present evidence that humans do differentiate CS+ and CS– faces even when controlling for the effect of exposure. Therefore, we believe that the major drive that leads to the separation of patterns along the specific axis are due to the affective nature of learning, rather than occurring merely as a result of exposure.

Eye-movements patterns can provide important insights about what the nervous system tries to achieve as they summarize the final outcome of complex interactions at the neuronal level (König et al., 2016). Our results demonstrate that changes induced by aversive generalization extend beyond autonomous systems or explicit subjective evaluations, but can also affect an entire sensory-motor loop at the systems level (Dowd et al., 2016). Furthermore, the methodology applied here can easily be extended to neuronal recordings, where gradients of activity during generalization have been successfully used to characterize selectivity of aversive representations. Therefore, it will be highly informative to test different hypotheses we outlined here using neuronal recordings with representational similarity analysis during the emergence of aversive representations.

## Materials and methods

### Participants

Participants were 74 naïve healthy males and females (n = 37 each) with normal (or corrected-to-normal) vision (age = 27 ± 4, M ± SD) and without history of psychiatric or neurological diseases, any medical condition or use of medication that would alter pain perception. Participants had not participated in any other study using facial stimuli in combination with aversive learning before. They were paid 12 Euros per hour for their participation in the experiment and provided written informed consent. All experimental procedures were approved by the Ethics committee of the General Medical Council Hamburg.

### Data sharing

The dataset used in this manuscript has been published as a dataset publication (Wilming et al., 2017). We publicly provide the stimuli as well as the Matlab (MathWorks, Natick MA) code necessary for the reproduction of all the results presented in this manuscript (Onat and Kampermann, 2017). The code can be used to download the data as well.

### Stimulus preparation and calibration of generalization gradient

Using a two-step procedure, we created a final set of 8 calibrated faces (Fig 1A, see also SFig 1) that were perceptually organized along a circular similarity continuum based on a model of the primary visual (V1) cortex. Using the FaceGen software (FaceGen Modeller 2.0, Singular Inversion, Ontario Canada) we created two gender-neutral facial identities and mixed these identities (0%/100% to 100%/0%) while simultaneously changing the gender parameters in two directions (more male or female). In the first step, we created a total of 160 faces by appropriately mixing the gender and identity parameters to form 5 concentric circles (see SFig 1) based on FaceGen defined parameter values for gender and identity. Using a simple model of the primary visual cortex known to closely mirror human perceptual similarity judgments (Yue et al., 2012), we computed V1 representations for each face after converting them to grayscale. The spatial frequency sensitivity of the V1 model was adjusted to match human contrast sensitivity function with bandpass characteristics between 1 and 12 cycles/degree, peaking at 6 cycles/degrees (Blakemore and Campbell, 1969). The V1 model consists of pair of Gabor filters in quadrature at five different spatial scales and eight orientations. The activity of these 40 channels were averaged in order to obtain one single V1 representation per face. We characterized the similarity relationship between the V1 representations of 160 faces using multidimensional scaling analysis with 2 dimensions (SFig 2). As expected, while two dimensions explained a large variance, the improvement with the addition of a third dimension was only minor, providing thus evidence that the physical properties of the faces were indeed organized along two-dimensions (stress values for 1D, 2D and 3D resulting from the MDS analysis were 0.42, .04, .03, respectively). The transformation between the coordinates of the FaceGen software values (gender and identity mixing values) and coordinates returned by the MDS analysis allowed us to gather FaceGen coordinates that would correspond to a perfect circle in the V1 model. In the second step, we thus generated 8 faces that corresponded to a perfect circle. This procedure ensured that faces used in this study were organized perfectly along a circular similarity continuum according to a simple model of primary visual cortex with well-defined bandpass characteristics known to mirror human similarity judgments. Furthermore it ensured that dimensions of gender and identity introduced independent variance on the faces.

To present these stimuli we resized them to 1000x1000 pixels (originals: 400x400) using bilinear interpolation, and slightly smoothed with a Gaussian kernel of 5 pixels with full-width at half maximum of 1.4 pixels to remove any possible pixel artifacts that could potentially lead participants to identify faces. Faces were then normalized to have equal luminance and root-mean-square contrast. The gray background was set to the same luminance level ensuring equal brightness throughout of the experiment. Faces were presented on a 20” monitor (1600 x 1200 pixels, 60 Hz) using Matlab R2013a (Mathworks, Natick MA) with psychophysics toolbox (Brainard, 1997; Pelli, 1997). The distance of the participants’ eyes to the stimulus presentation screen was 50 cm. The center of the screen was at the same level as the participants’ eyes. Faces spanned horizontally ∼17° and vertically ∼30°, aiming to mimic a typical face-to-face social situation. Stimuli are available in (Onat and Kampermann, 2017).

### Experimental paradigm

The fear conditioning paradigm (similar to (Onat and Büchel, 2015)) consisted of baseline, conditioning and test (or generalization) phases (Fig 2A). Participants were instructed that the delivery of UCSs during baseline would not be associated with faces, however in the following conditioning and generalization phases they were instructed that shocks would be delivered after particular faces have been presented. In all three phases, subjects were instructed to press a button when an oddball stimulus appeared on the screen.

Four equivalent runs with exactly same number of trials were used during baseline (1 run) and generalization phases (3 runs) consisting of 120 trials per run (∼10 minutes). Every run started with an eye-tracker calibration. Between runs participants took a break and continued with the next run in a self-paced manner. We avoided having more than 1 run in the baseline period in order not to induce fatigue in participants. At each run during the baseline and generalization phases, 8 faces were repeated 11 times, UCS trials occurred 5 times and one oddball was presented. This consisted of a blurred unrecognizable face, which volunteers were instructed to press a key. We presented 26 null trials with no face presentation but otherwise the same trial structure (see below sequence optimization). In order to keep arousal levels comparable to the generalization phase, UCSs were also delivered during baseline, however they were fully predictable by a shock symbol therefore avoiding any face to UCS associations.

During the conditioning phase, participants saw only the CS+ and the CS– faces (and null trials). These consisted of 2 maximally dissimilar faces separated by 180° on the circular similarity continuum and randomly assigned for every participant in a balanced manner. The conditioning was 124 trials long (∼10 minutes) and CS+ and CS– faces were repeated 25 times. CS+ faces were additionally presented 11 times with the UCSs, resulting in a reinforcement rate of ∼30 %. The same reinforcement ratio was used during the subsequent generalization phase in order to avoid extinction of the learnt associations.

Stimulus presentation sequence was optimized for the deconvolution of the skin-conductance responses. This regards the choice of conditions and the duration of interstimulus intervals. Faces were presented using a rapid-event design with a stimulus onset asynchrony of 6 seconds and stimulus duration of 1.5 seconds. The presentation sequence was optimized using a modified m-sequence with 11 different conditions (Buracas and Boynton, 2002; Liu and Frank, 2004) (8 faces, UCS, oddball, null). An m-sequence is preferred as it balances all transitions from condition *n* to *m* (thus making the sequence as unpredictable as possible for the participant) while providing an optimal design efficiency (thus making deconvolution of autonomic skin conductance responses more reliable). However, all conditions in an m-sequences appear equally number of times. Therefore, in order to achieve the required reinforcement ratio (∼30%), we randomly pruned UCS trials and transformed them to null trials. Similarly, oddball trials were pruned to have an overall rate of ∼1%. This resulted in a total of 26 null trials. While this deteriorated the efficiency of the m-sequence, it was still a good compromise as the resulting sequence was much more efficient than a random sequence. Resulting from the intermittent null trials, SAOs were 6 or 12 seconds approximately exponentially distributed.

Face onsets were preceded by a fixation-cross, which appeared randomly outside of the face either on the left or right side along an virtual circle (r = 19.6°, +/- 15° above and below the horizontal center of the image). The side of fixation-cross was balanced across conditions to avoid confounds that might occur (Arizpe et al., 2015). Therefore, the first fixation consisted of a landing fixation on the face.

### Calibration and delivery of electric stimulation

Mild electric shocks were delivered by a direct current stimulator (Digitimer Constant Current Stimulator, Hertfordshire UK), applied by a concentric electrode (WASP type, Speciality Developments, Kent UK) that was firmly connected to the back of the right hand and fixated by a rubber glove to ensure constant contact with the skin. Shocks were trains of 5-ms pulses at 66Hz, with a total duration of 100 ms. During the experiment, they were delivered right before the offset of the face stimulus. The intensity of the electric shock applied during the experiment was calibrated for each participant before the start of the experiment. Participants underwent a QUEST procedure (Watson and Pelli, 1983) presenting UCSs with varying amplitudes selected by an adaptive algorithm and were required to report whether a given trial was “painful” or “not painful” in a binary fashion using a sliding bar. The QUEST procedure was repeated twice to account for sensitization/habituation effects, thus obtaining a reliable estimate. Each session consisted of 12 stimuli, starting at an amplitude of 1mA. The subjective pain threshold was the intensity that participants would rate as “painful” with a probability of 50%. The amplitude used during the experiment was 2 times this threshold value. Before starting the actual experiment, participants were asked to confirm whether the resulting intensity was bearable. If not then the amplitude was incrementally reduced and the final amplitude was used for the rest of the experiment.

### Eye tracking and fixation density maps

Eye tracking was done using an Eyelink 1000 Desktop Mount system (SR Research, Ontario Canada) recording the right eye at 1000 Hz. Participants placed their head on a headrest supported under the chin and forehead to keep a stable position. Participants underwent a 13 point calibration / validation procedure at the beginning of each run (1 Baseline run, 1 Conditioning run and 3 runs of Generalization). The average mean-calibration error across all runs was Mean = 0.36°, Median = .34°, SD = 0.11. 91% of all runs had a calibration better than or equal to .5°.

Fixation events were identified using commonly used parameter definitions (Wilming et al., 2017) (Eyelink cognitive configuration: saccade velocity threshold = 30° / second, saccade acceleration threshold = 8000° per second^2^, motion threshold = .1°). Fixation density maps (FDMs) were computed by spatially smoothing (Gaussian kernel of 1° of full width at half maximum) a 2D histogram of fixation locations, and were transformed to probability densities by normalizing to unit sum. FDMs included the center 500x500 pixels, including all facial elements where fixations were mostly concentrated (∼95% of all fixations).

### Shock expectancy ratings and autonomic recordings

After baseline, conditioning and generalization phases, participants rated different faces for subjective shock expectancy by answering the following question, “*How likely is it to receive a shock for this face?*”. Faces were presented in a random order and rated twice. Subjects answered using a 10 steps scale ranging from “*very unlikely*” to “*very likely*” and confirmed by a button press in a self-paced manner.

Electrodermal activity evoked by individual faces was recorded throughout the three phases. Reusable Ag/AgCl electrodes filled with isotonic gel were connected to the palm of the subject’s left hand using adhesive collars, placed in thenar/hypothenar configuration. Skin-conductance responses were continuously recorded using a Biopac MP100 AD converter and amplifier system at a sampling rate of 500 Hz. Using the Ledalab toolbox (Benedek and Kaernbach, 2010a, 2010b), we decomposed the raw data to phasic and tonic response components after downsampling it to 100 Hz. Ledalab applies a positively constrained deconvolution technique in order to obtain phasic responses for each single trial. We averaged single-trial phasic responses separately for each condition and experimental phase to obtained 21 average values (9 (8 faces + 1 null condition) from baseline and generalization and 3 (2 faces + 1 null condition) from the conditioning phase). CS+ trials with UCS were excluded from this analysis. These values were first log-transformed (log_10_(1+SCR)) and subsequently z-scored for every subject separately (across all conditions and phases), then averaged across subjects. Therefore, negative values indicate phasic responses that are smaller than the average responses recorded throughout the experiment. Due to technical problems, SCR data could only be analyzed for n = 63 out of the 74 participants.

### Nonlinear modelling and model comparison

We fitted a von Mises function (circular Gaussian) to generalization profiles obtained from subjective ratings, skin-conductance responses and fixation counts at different ROIs by minimizing the following likelihood term in (1) following an initial grid-search for parameters

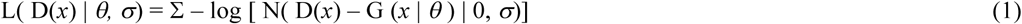

where *x* represents signed angular distances from a given volunteer’s CS+ face; G(*x*|*θ*) is a von Mises-like function that was used to model the adversity tuning. It is defined by the parameter vector *θ,* which codes for the amplitude (difference between peak and base), location (peak position), precision and offset (base value) of the resulting generalization profile; D(*x*) represents the observed generalization profile for different angular distances; and N(x| 0, *σ*) is the normal probability density function with mean zero and standard deviation of *σ*. The fitting procedure consisted of finding parameters values that minimized the sum of negative log-transformed probability values. Using log-likelihood ratio test we tested whether this model performed better than a null model consisting of a horizontal line, effectively testing the significance of the additional variance explained by the model. G(x) was a scaled and shifted version of a normalized von Mises function in the form

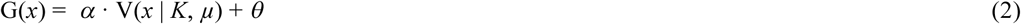

*α* represents the depth of adversity tuning which corresponds to the difference between peak and baseline responses, and *θ* sets to the baseline level. *K* and *µ* controls the precision of the tuning and the peak position of adversity tuning, respectively. V(x) is a modified von Mises function that is scaled to fit between 0 and 1 using the following equation:

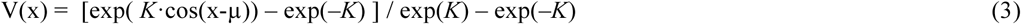

### Fixation-pattern similarity analysis

FPSA was conducted on single participants. Condition specific FDMs (8 faces per baseline and generalization phases) were computed by collecting all fixations across trials on a single map which was then normalized to unit sum. We corrected FDMs by removing the common mean pattern (done separately for baseline and generalization phases). We used 1 - Pearson correlation as the similarity metric. This resulted in a 16x16 similarity matrix per subject. Statistical tests for element-wise comparison of the similarity values were conducted after Fisher transformation of correlation values. The multidimensional scaling was conducted on the baseline and generalization phases jointly using the 16x16 similarity matrix as input (*mdscale* in MATLAB). Importantly, as the similarity metric is extremely sensitive to the signal to noise ratio (Diedrichsen et al., 2011) present in the FDMs, we took precautions that the number of trials between generalization and baseline phases were exactly the same in order to avoid differences that would have been caused by different signal to noise ratios. To account for unequal number of trials during the baseline (11 repetitions) and generalization (3 runs x 11 = 33 repetitions) phases, we computed a similarity matrix for each run separately in the generalization phase. These were later averaged across runs for a given participant. This ensured that FDMs of the baseline and generalization phases had comparable signal-to-noise ratios, therefore not favoring the generalization phase for having more trials.

We generated 3 different models based on a quadrature decomposition of a circular similarity matrix. A circular similarity matrix of 8x8 can be obtained using the term **M**⊗**M**, where **M** is a 8x2 matrix in form of [cos(*x*) sin(*x*)], and the operator ⊗ denotes the outer product. *x* represents angular distances from the CS+ face, is equal to 0 for CS+ and π for CS–. Therefore, while cos(*x*) is symmetric around the CS+ face, sin(*x*) is shifted by 90°. For the Bottom-up Saliency and Increased Arousal models (Fig 1B and C) we used M⊗M as a predictor together with a constant intercept. For the tuned exploration model depicted in Fig 1D, we used cos(*x*) ⊗cos(*x*) and sin(*x*)⊗sin(*x*) to independently model ellipsoid expansion along the specific and unspecific directions, respectively. Together with the intercept this model comprised 3 predictors. Finally the aversive generalization model (Fig 1E) was created using the predictors of the tuned exploration model in conjunction with a two-dimensional Gaussian centered on the CS+ face (in total 4 predictors). We tested different widths for the Gaussian and took the one that resulted in the best fit. This was equal to 65° of FWHM and similar to the values we observed for univariate explicit ratings and SCR responses.

All linear modeling was conducted using non-redundant, vectorized forms of the symmetric dissimilarity matrices. For a 8x8 dissimilarity matrix this resulted in a vector of 28 entries. Different models were fitted as mixed-effects, where intercept and slope contributed both as fixed- and random-effects (*fitlme* in Matlab). We selected mixed-effect models as these performed better than models defined uniquely with fixed-effects on intercept and slope. To do model selection, we used Bayesian information criterion (BIC) as it compensates for an increase in the number of predictors between different models. Additionally, different models were also fitted to single participants (*fitlm* in Matlab) and the parameter estimates were separately tested for significance using *t-test*.

For the analysis of temporal and spatial unfolding of adversity specific exploration patterns, the same analysis was run, but restricted to include only the given time windows / fixations. In this analysis, we only included participants who had a significant fear-tuning of explicit shock ratings from the generalization phase (n = 61). For the time-windowed approach, periods of 500 ms were used, repeated shifted by 50 ms, thereby obtaining a rolling-window analysis, while fixation-wise analyses were based on FDMs that included only on the given (1^st^, 2^nd^, and so on) fixation.

## Acknowledgements

The authors wish to thank Tim Kietzmann for his input on an early version of this manuscript, Cliodhna Quigley for proof-reading, Helen Blank for comments, Patricia Billaudelle and Katrin Harland for their assistance with data collection. Selim Onat is supported by the DFG SFB TRR 58.

## Author contributions

Conceptualization: SO, CB

Data Curation: SO

Formal Analysis: LK, NW, SO

Funding Acquisition: CB

Investigation: LK, SO

Methodology: LK, NW, SO

Project Administration: SO

Resources: CB

Software: LK, SO

Supervision: SO

Validation: LK, NW, AA, CB, SO

Visualization: LK, SO

Writing - Original Draft Preparation: LK, SO

Writing – Review, Editing: LK, NW, AA, SO

## Supporting Material

**SFig 1.**
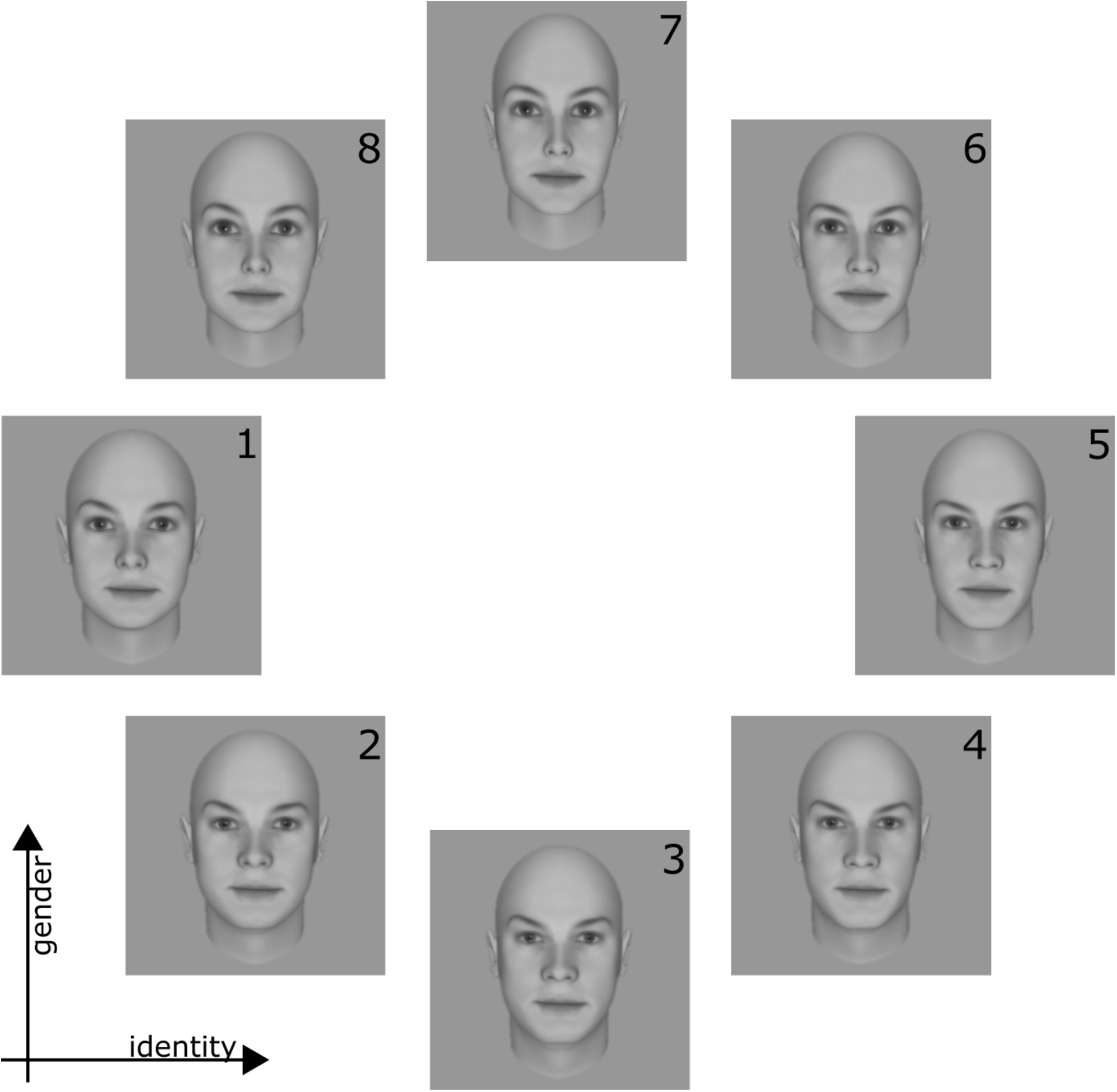
Face Stimuli. Set of 8 faces that were calibrated to form a circular similarity continuum. Faces vary along the two dimensions of gender (vertical axis) and identity (horizontal axis). See SFig 2 for the calibration process.

**SFig 2.**
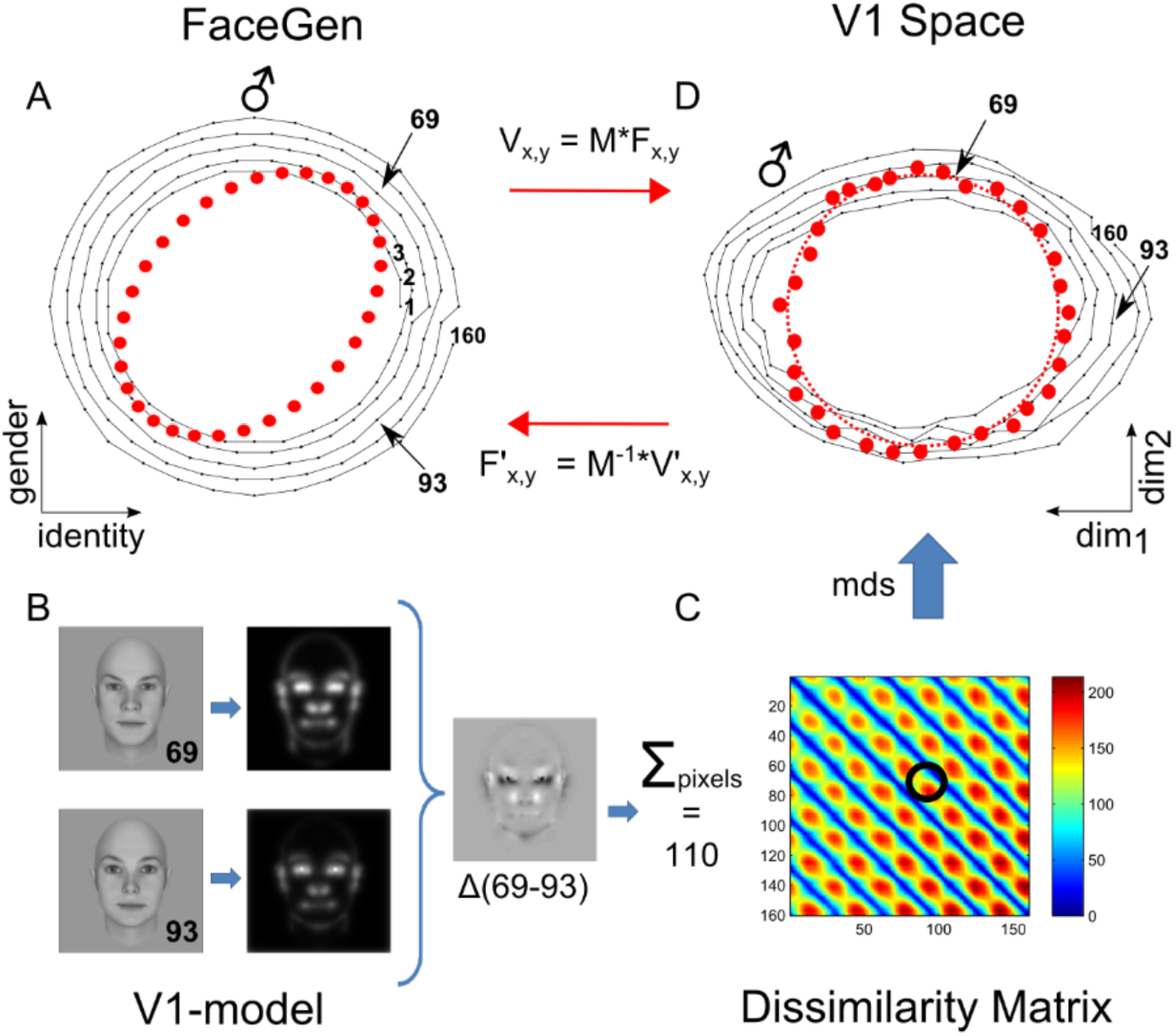
Calibration of faces using a simple V1 model tuned to human psychophysics. (A) Using the FaceGen software, 160 faces forming five concentric circles were generated with coordinates varying in gender and identity dimensions (connected black dots in the left panel). Maximally male faces are located at 12 o’clock direction and indicated with the male symbol. (B) V1 representations of faces were modelled according to (Yue et al., 2012). This is illustrated for faces 69 and 93. The difference between these two faces resulted in a Euclidean distance of 110. The pair-wise Euclidean distance for all the 160 faces are shown in (C) as a dissimilarity matrix. The resulting dissimilarity matrix exhibits 5 major bands corresponding to 5 concentric circles. By applying MDS, we obtained the representational space of V1 shown in (A, right panel). Note that the most male face is 45° counter-clockwise rotated with respect to the main axes of in V1 representation. The mapping between FaceGen coordinates and V1 representational space thus involved a rotation and scaling which was captured by the matrix *M*. We therefore used the inverse of *M*, to achieve coordinates of perfect circularity based on this V1 model. This ensured that faces along the similarity continuum were characterized by controlled changes for every angular step based on the model used.

**SFig 3.**
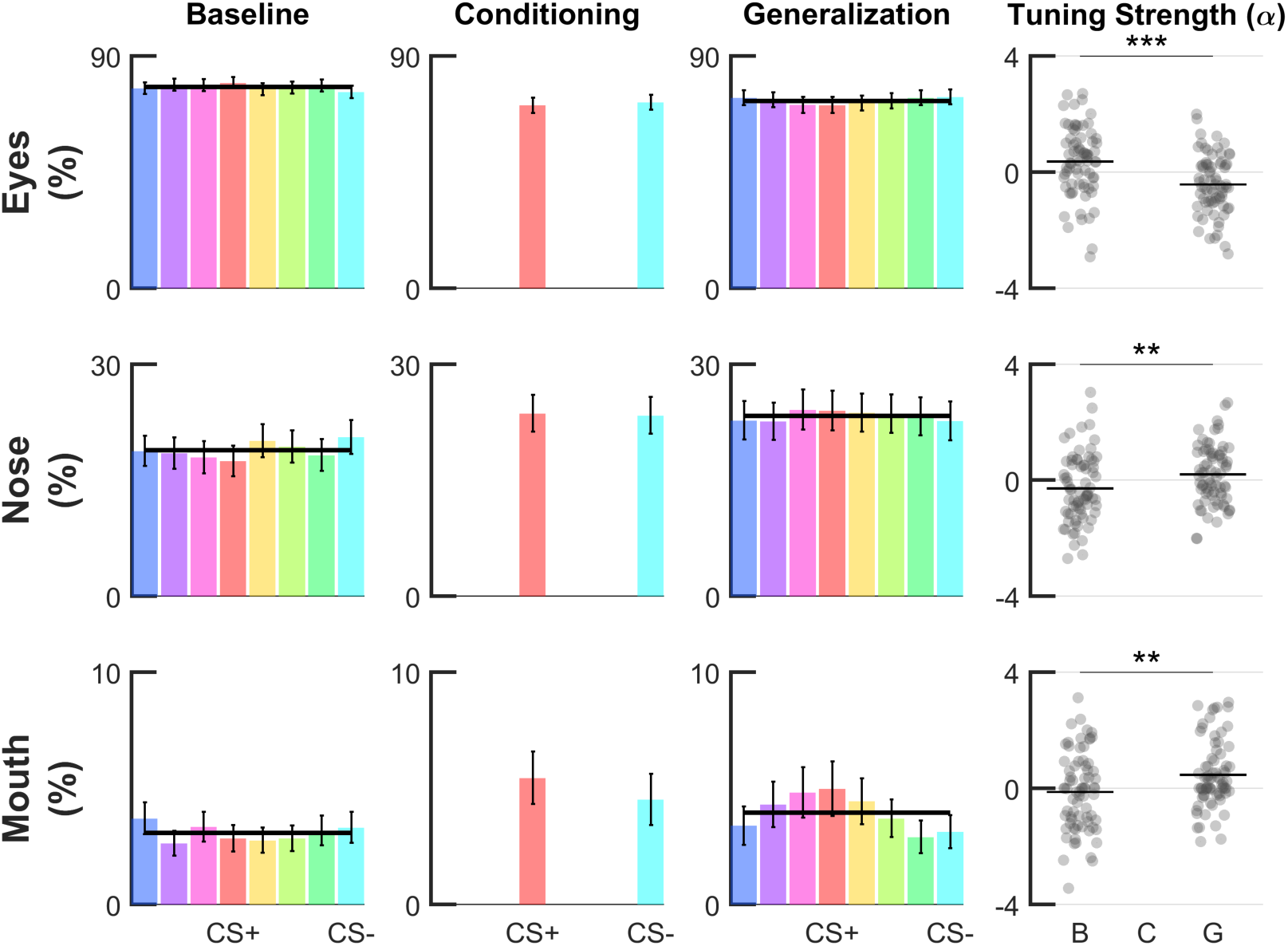
Generalization profiles on ROI-based fixation counts. Group-level fear-tuning based on percentage of fixations in a given ROI (n = 74) for different phases. Fixation count data expressed as percentages are based on three major facial regions, i.e. eyes, nose and mouth (see Fig. 4). Responses are aligned to the CS+ of each volunteer separately (errorbars: SEM across subjects). Black horizontal lines or curves indicate the winning model (p < .001, log-likelihood ratio test), i.e. horizontal null model or the Gaussian model. In all situations, the null horizontal model was favored by the model comparison procedure. Scatter plots show amplitude parameter α of single-subject Gaussian fits. Horizontal lines within the scatterplots depict group-level means, asterisks indicate significant differences in α (compared to baseline phase, paired t-test, ***: p < .001, **: p < .01).

**S1 Table.**
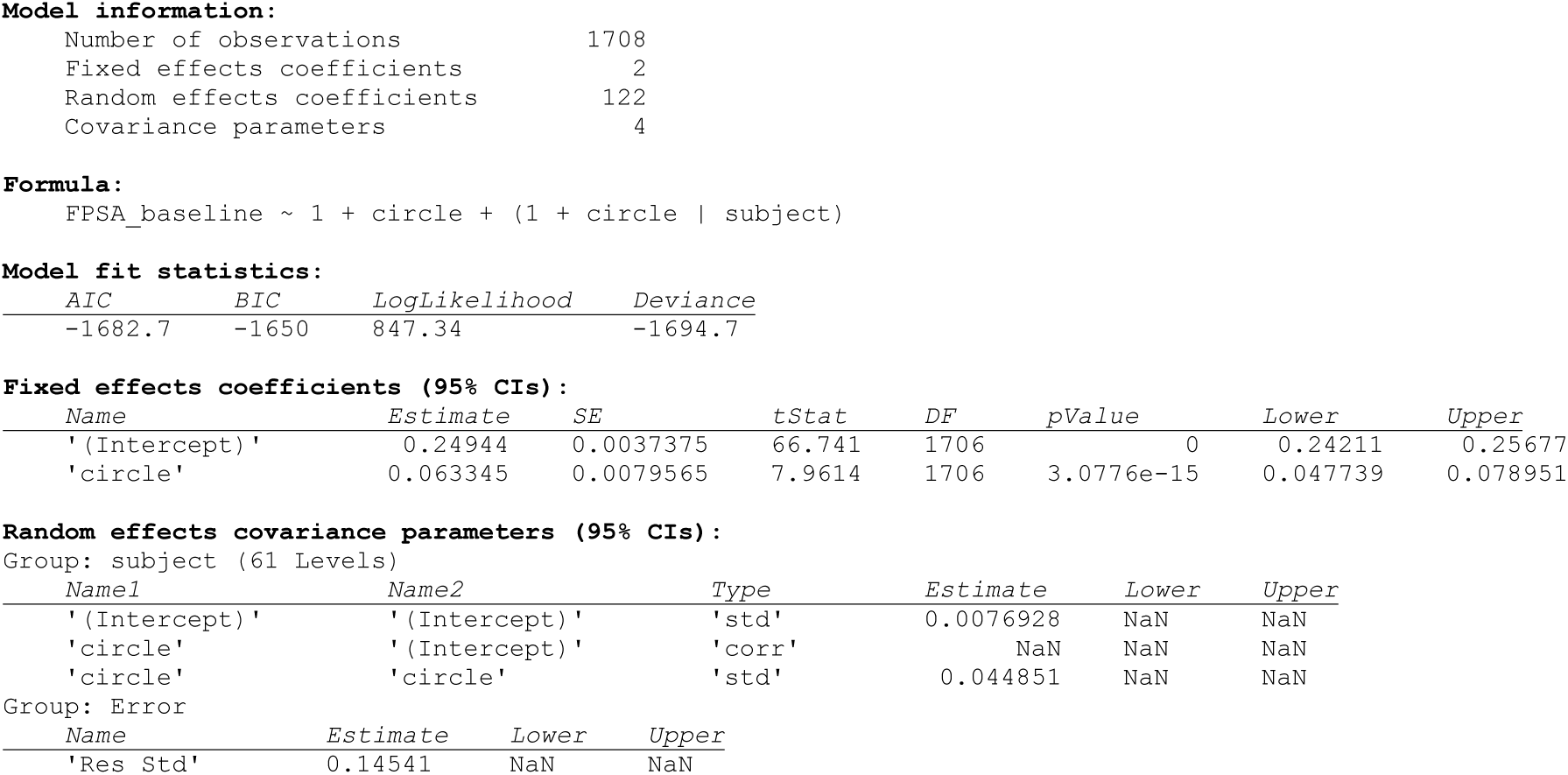
Mixed-effects modeling of the similarity matrices during **the baseline phase with the Perceptual Similarity model** shown in Fig 1B.

**S2 Table.**
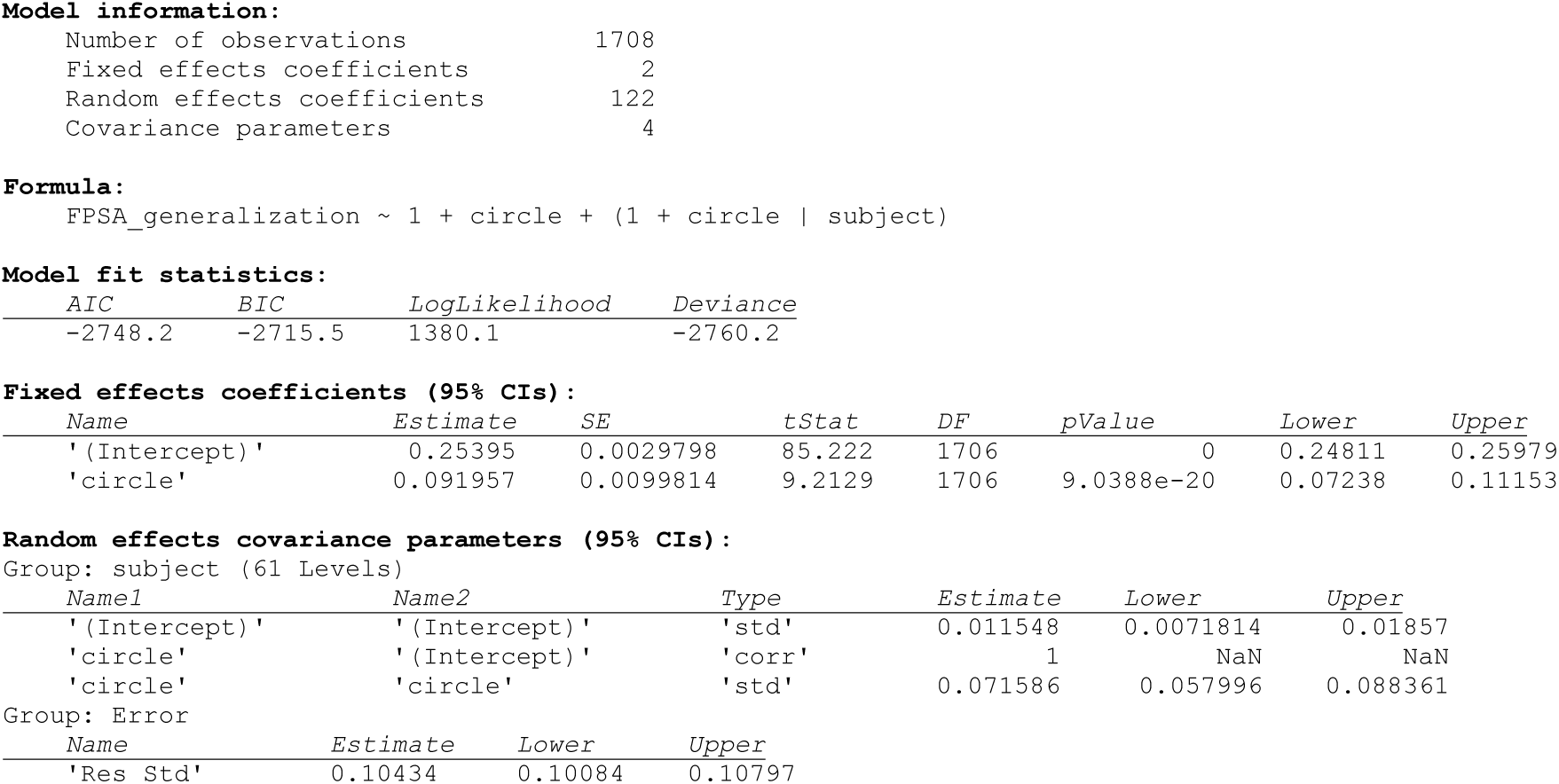
Mixed-effects modeling of the similarity matrices during **the generalization phase with the Perceptual Similarity model** shown in Fig 1C.

**S3 Table.** Mixed-effects modeling of the similarity matrices during **the baseline phase with the Adversity Categorization model** shown in Fig 1D.

|  |  |
| --- | --- |
| Number of observations | 1708 |
| Fixed effects coefficients | 3 |
| Random effects coefficients | 183 |
| Covariance parameters | 7 |

| AIC | BIC | LogLikelihood | Deviance |
| --- | --- | --- | --- |
| -1764 | -1709.6 | 892.01 | -1784 |

Fixed effects coefficients (95% CIs):
| Name | Estimate | SE | tStat | DF | pValue | Lower | Upper |
| --- | --- | --- | --- | --- | --- | --- | --- |
| '(Intercept)' | 0.24944 | 0.0035811 | 69.654 | 1705 | 0 | 0.24242 | 0.25646 |
| 'specific' | 0.057755 | 0.013652 | 4.2304 | 1705 | 2.457e-05 | 0.030977 | 0.084532 |
| 'unspecific' | 0.068935 | 0.010663 | 6.4647 | 1705 | 1.3216e-10 | 0.048021 | 0.08985 |

Random effects covariance parameters (95% CIs): Group: subject (61 Levels)
| Name1 | Name2 | Type | Estimate | Lower | Upper |
| --- | --- | --- | --- | --- | --- |
| '(Intercept)' | '(Intercept)' | 'std' | 0.0081486 | 0.0035081 | 0.018928 |
| 'specific' | '(Intercept)' | 'corr' | 0.81502 | 0.80893 | 0.82093 |
| 'unspecific' | '(Intercept)' | 'corr' | 0.30346 | -0.0053757 | 0.55945 |
| 'specific' | 'specific' | 'std' | 0.0913 | 0.071842 | 0.11603 |
| 'unspecific' | 'specific' | 'corr' | -0.30479 | -0.56025 | 0.0036179 |
| 'unspecific' | 'unspecific' | 'std' | 0.062467 | 0.045811 | 0.085177 |

Group: Error
| Name | Estimate | Lower | Upper |
| --- | --- | --- | --- |
| 'Res Std' | 0.13817 | 0.13344 | 0.14306 |

**S4 Table.** Mixed-effects modeling of the similarity matrices during **the generalization phase with the Adversity Categorization** model shown in Fig 1D.

| AIC | BIC | LogLikelihood | Deviance |
| --- | --- | --- | --- |
| -2951.7 | -2897.3 | 1485.9 | -2971.7 |

Fixed effects coefficients (95% CIs):
| Name | Estimate | SE | tStat | DF | pValue | Lower | Upper |
| --- | --- | --- | --- | --- | --- | --- | --- |
| '(Intercept)' | 0.25395 | 0.0028004 | 90.683 | 1705 | 0 | 0.24846 | 0.25944 |
| 'specific' | 0.1205 | 0.014472 | 8.3259 | 1705 | 1.6951e-16 | 0.09211 | 0.14888 |
| 'unspecific' | 0.063418 | 0.010001 | 6.3412 | 1705 | 2.9123e-10 | 0.043803 | 0.083034 |

Random effects covariance parameters (95% CIs): Group: subject (61 Levels)
| Name1 | Name2 | Type | Estimate | Lower | Upper |
| --- | --- | --- | --- | --- | --- |
| '(Intercept)' | '(Intercept)' | 'std' | 0.011778 | 0.0077913 | 0.017805 |
| 'specific' | '(Intercept)' | 'corr' | 0.91907 | NaN | NaN |
| 'unspecific' | '(Intercept)' | 'corr' | 0.6935 | NaN | NaN |
| 'specific' | 'specific' | 'std' | 0.10647 | 0.087673 | 0.12931 |
| 'unspecific' | 'specific' | 'corr' | 0.35345 | 0.34746 | 0.35941 |
| 'unspecific' | 'unspecific' | 'std' | 0.068277 | 0.054449 | 0.085617 |

Group: Error
| Name | Estimate | Lower | Upper |
| --- | --- | --- | --- |
| 'Res Std' | 0.09517 | 0.091916 | 0.09854 |

**S5 Table.** Mixed-effects modeling of the similarity matrices during **the baseline phase with the Adversity Tuning model** shown in Fig 1E.

**Model information:**
|  |  |
| --- | --- |
| Number of observations | 1708 |
| Fixed effects coefficients | 4 |
| Random effects coefficients | 244 |
| Covariance parameters | 11 |

**Model fit statistics:**
| AIC | BIC | LogLikelihood | Deviance |
| --- | --- | --- | --- |
| -1755.5 | -1673.9 | 892.76 | -1785.5 |

**Fixed effects coefficients (95% CIs):**
| Name | Estimate | SE | tStat | DF | pValue | Lower | Upper |
| --- | --- | --- | --- | --- | --- | --- | --- |
| '(Intercept)' | 0.24181 | 0.046873 | 5.1589 | 1704 | 2.7749e-07 | 0.14988 | 0.33374 |
| 'specific' | 0.057483 | 0.013761 | 4.1774 | 1704 | 3.0984e-05 | 0.030494 | 0.084473 |
| 'unspecific' | 0.069013 | 0.010773 | 6.4063 | 1704 | 1.9241e-10 | 0.047884 | 0.090142 |
| 'Gaussian' | 0.015884 | 0.097484 | 0.16294 | 1704 | 0.87058 | -0.17532 | 0.20709 |

**Random effects covariance parameters (95% CIs):** Group: subject (61 Levels)
| Name1 | Name2 | Type | Estimate | Lower | Upper |
| --- | --- | --- | --- | --- | --- |
| '(Intercept)' | '(Intercept)' | 'std' | 0.054362 | 0.022712 | 0.13012 |
| 'specific' | '(Intercept)' | 'corr' | 0.16475 | NaN | NaN |
| 'unspecific' | '(Intercept)' | 'corr' | -0.98746 | -0.9876 | -0.98731 |
| 'Gaussian' | '(Intercept)' | 'corr' | -0.99227 | -0.99237 | -0.99216 |
| 'specific' | 'specific' | 'std' | 0.091417 | 0.07191 | 0.11622 |
| 'unspecific' | 'specific' | 'corr' | -0.31842 | -0.32004 | -0.31681 |
| 'Gaussian' | 'specific' | 'corr' | -0.041041 | NaN | NaN |
| 'unspecific' | 'unspecific' | 'std' | 0.063537 | 0.046652 | 0.086532 |
| 'Gaussian' | 'unspecific' | 'corr' | 0.96022 | NaN | NaN |
| 'Gaussian' | 'Gaussian' | 'std' | 0.12196 | 0.054259 | 0.27414 |

Group: Error
| Name | Estimate | Lower | Upper |
| --- | --- | --- | --- |
| 'Res Std' | 0.13805 | 0.13333 | 0.14293 |

**S6 Table.** Mixed-effects modeling of the similarity matrices during **the generalization phase with the Adversity Tuning model** shown in Fig 1E.

**Model fit statistics:**
| AIC | BIC | LogLikelihood | Deviance |
| --- | --- | --- | --- |
| -2945.9 | -2864.2 | 1487.9 | -2975.9 |

**Fixed effects coefficients (95% CIs):**
| Name | Estimate | SE | tStat | DF | pValue | Lower | Upper |
| --- | --- | --- | --- | --- | --- | --- | --- |
| '(Intercept)' | 0.22 | 0.032419 | 6.786 | 1704 | 1.5856e-11 | 0.15641 | 0.28359 |
| 'specific' | 0.11929 | 0.014301 | 8.3414 | 1704 | 1.4959e-16 | 0.091239 | 0.14734 |
| 'unspecific' | 0.063765 | 0.010056 | 6.3412 | 1704 | 2.913e-10 | 0.044042 | 0.083488 |
| 'Gaussian' | 0.070672 | 0.067933 | 1.0403 | 1704 | 0.29834 | -0.062568 | 0.20391 |

**Random effects covariance parameters (95% CIs):** Group: subject (61 Levels)
| Name1 | Name2 | Type | Estimate | Lower | Upper |
| --- | --- | --- | --- | --- | --- |
| '(Intercept)' | '(Intercept)' | 'std' | 0.044693 | 0.010711 | 0.18649 |
| 'specific' | '(Intercept)' | 'corr' | -0.89633 | -0.89679 | -0.89586 |
| 'unspecific' | '(Intercept)' | 'corr' | -0.73005 | -0.73122 | -0.72888 |
| 'Gaussian' | '(Intercept)' | 'corr' | -0.99995 | -0.99996 | -0.99995 |
| 'specific' | 'specific' | 'std' | 0.1047 | 0.08608 | 0.12734 |
| 'unspecific' | 'specific' | 'corr' | 0.35136 | 0.35001 | 0.3527 |
| 'Gaussian' | 'specific' | 'corr' | 0.9005 | 0.90014 | 0.90086 |
| 'unspecific' | 'unspecific' | 'std' | 0.068748 | 0.054885 | 0.086113 |
| 'Gaussian' | 'unspecific' | 'corr' | 0.72353 | 0.7229 | 0.72415 |
| 'Gaussian' | 'Gaussian' | 'std' | 0.11757 | 0.03792 | 0.36454 |

Group: Error
| Name | Estimate | Lower | Upper |
| --- | --- | --- | --- |
| 'Res Std' | 0.095034 | 0.091784 | 0.098399 |

